# Molecular Characterization of invasive *Enterobacteriaceae* from Pediatric Patients in Central and Northwestern Nigeria

**DOI:** 10.1101/2020.02.21.959338

**Authors:** Carissa Duru, Grace M Olanipekun, Vivian Odili, Nicholas J Kocmich, Amy Rezac, Theresa O Ajose, Nubwa Medugu, Bernard Ebruke, Charles Esimone, Stephen Obaro

## Abstract

**Background:** Bacteremia is a leading cause of death in developing countries but etiologic evaluation is infrequent and empiric antibiotics are not evidence-based. Very little is known about the types of extended-spectrum β-lactamases (ESBL) in pediatric bacteremia patients in Nigeria. We evaluated the patterns of ESBL resistance in children enrolled into surveillance for community acquired bacteremic syndromes across health facilities in Central and Northwestern Nigeria.

**Method:** Blood culture from suspected cases of sepsis from children age less than 5 years were processed using automated Bactec^®^ incubator System from Sept 2008-Dec 2016. *Enterobacteriacea* were identified to the species level using Analytical Profile Index (API20E^®^) identification strip and antibiotic susceptibility profile was determined by the disc diffusion method. The multidrug resistant strains were then screened and confirmed for extended spectrum beta lactamase (ESBL) production by the combination disc method as recommended by Clinical and Laboratory Standard Institute (CLSI). Real time PCR was used to elucidate the genes responsible for ESBL production characterize the resistance genes

**Result:** Of 21,000 children screened from Sept 2008-Dec 2016, 2,625(12.5%) were culture-positive. A total of 413 *Enterobacteriaceae* available for analysis were screened for ESBL. ESBL production was detected in 160/413(38.7%), comprising *Klebsiella pneumoniae* 105/160(65.6%), *Enterobacter cloacae* 21/160(13.1%), *Escherichia coli* 22/160(13.8%), *Serratia* species 4/160(2.5%), *Pantoea* species 7/160(4.4%) and *Citrobacter* species 1/160(0.6%). Of the 160 ESBL-producing isolates, high resistance rates were observed among ESBL-positive isolates for Ceftriaxone (92.3%), Aztreonam (96.8%), Cefpodoxime (96.25%), Cefotaxime (98.75%) and sulphamethoxazole-trimethoprim (90%), while 87.5 %, 90.63%, and 91.87% of the isolates were susceptible to Imipenem, Amikacin and Meropenem respectively. Frequently detected resistance genes were *bla*TEM 83.75%) (134/160), and, *bla*CTX-M 83.12% (133/160) followed by *bla*SHV genes 66.25% (106/160). Co-existence of *bla*CTX-M, *bla*TEM and *bla*SHV was seen in 94/160 (58.8%), *bla*CTX-M and *bla*TEM in 118/160 (73.8%), *bla*TEM and *bla*SHV in 97/160 (60.6%) and *bla*CTX-M and *bla*SHV in 100/160 (62.5%) of isolates tested.

**Conclusion:** Our results indicate a high prevalence of ESBL resistance to commonly used antibiotics in *Enterobacteriaceae* isolates from bloodstream infections in children in this study. Careful choice of antibiotic treatment options and further studies to evaluate transmission dynamics of resistance genes could help in the reduction of ESBL resistance in these settings.

## Introduction

Bacterial blood stream infection (BSI) is a major public-health concern especially in developing countries as it is one of the leading causes of death. In Nigeria as in most developing countries, sub optimal laboratory methods contribute to improper diagnosis of BSI in children [1]. The absence of etiologic diagnosis and clinical microbiology laboratories implies that patients are often treated with broad spectrum antibiotics. This unguided approach to clinical care promotes the incidence of drug resistant bacteria.

Antibiotic resistance has become a concern worldwide and in *Enterobacteriaceae*, the production of β-lactamase remains the most important mediator of β-lactam resistance [2]. Extended-spectrumβ-lactamases (ESBLs) are a rapidly evolving group of β-lactamases which hydrolyze the extended-spectrum cephalosporins, the penicillins, as well as aztreonam, but not carbapenems [3]. These ESBL-producing bacteria are also resistant to other class of antimicrobial agents such as aminoglycosides, trimethoprim/sulfamethoxazole, and quinolones thereby leading to Multidrug-resistance (MDR), which often complicate the treatment of severe bacterial infections [2,4,5]. The World Health Organization (WHO) has declared infections caused by multidrug resistant bacteria as an emerging global health problem of major public health concern [6].

The ESBL enzymes were initially recognized in clinical isolates in the 1980s; they are derived mainly from the TEM or SHV types of β-lactamases, by point mutations in the parent enzymes which did not possess extended-spectrum β-lactam substrate activity [3, 7].

More than 200 ESBLs have been identified so far, apart from the TEM, SHV and CTX-M types, other clinically relevant types of ESBLs include the VEB, PER, GES, TLA, IBC, SFO-1, BES-1 and BEL-1 types [7].

The prevalence of ESBL-producing bacteria has been reported worldwide [8–13], while there are a number of publications on ESBL-producing bacteria causing clinical infections [14–19] in Nigeria, report on the characterization of invasive isolates from infants and children is scanty and since antimicrobial resistance varies greatly among geographical settings, it is crucial to base empiric therapy of severe infections such as BSI on comprehensive knowledge of the prevalence and antimicrobial resistance patterns of locally isolated bacteria.

Therefore, the aim of this study was to investigate the genetic composition of ESBL-E from pediatric BSI patients.

## Methods

### Isolate collection

The study was conducted at 7 hospitals in Federal Capital Territory (FCT) and 3 in Kano Nigeria as previously described from Sept 2008 - Dec 2016 [1, 20]. Children aged less than five years were enrolled at the different sites, blood specimens of 1-3 ml were collected using the vacutainer set, after aseptically cleansing the skin with alcohol swab and povidone-iodine, the specimen was collected directly into an aerobic blood culture bottle (BD BactecPeds Plus/F culture vials; Becton Dickinson, Ireland), and incubated in an automated Bactec 9050 machine. All positive bottles were sub cultured onto MacConkey, Sheep blood and chocolate agar plates at 37°C for 24 h.

A total number of 887 culture-positive *Enterobacteriaceae* were obtained. Of the 887 isolates, 474 salmonella species which have been reported by Obaro et al. 2015 were excluded [20], therefore 413 including *Escherichia coli, Klebsiella* species, *Enterobacter* species, *Serratia marcescens, Pantoea* species, *Salmonella Typhi* and *Citrobacter* species from September 2008 to December 2016 were included. The isolates were stored in 10% skim milk glycerol at −80 °C.

### Phenotypic screening and confirmation of ESBL

*Enterobacteriaceae* were identified to the species level using Analytical Profile Index (API 20E) identification strip (Biomeriux Inc, France). Phenotypic ESBL production was confirmed with the combination disc diffusion test with clavulanic acid. CLSI confirmatory test was considered positive when the inhibition zone produced by the discs in combination clavulanate increased ≥5 mm than the disks without the clavulanate (CLSI, 2017), [21].

Susceptibility was determined using the disk diffusion method on Mueller Hinton agar as recommended by the Clinical and Laboratory Standard Institute (CLSI) (CLSI, 2017), (21). Susceptibility was tested against Amoxicillin/clavunate (20/10μg), Cefoxitin (30μg), Trimethoprim-Sulfamethoxazole (1.25/23.75μg), Ciprofloxacin (5μg), Ceftriaxone (30μg), Amikacin (30μg), Cefpodoxime (10μg), ceftazidime (30μg), Imipenem (10μg) and Meropenem (10μg), Cefotaxime (30μg), Piperacillin tazobactam (110μg), Aztreonam (30μg) and Tigecycline (15μg), (Oxoid Ltd, Basingstoke, Hampshire, England) and interpretation of breakpoint was according to CLSI, 2017 guidelines.

### Molecular identification of ESBL genes

All phenotypic ESBL producers were analyzed by Real time PCR for the presence of genes encoding TEM, SHV, CTX-M, Primers and Probes were designed for ESBL producing genes by LGC, Biosearch, USA based on primers used by Roschanski *et al*., 2014 [22].

Genomic DNA was extracted using Maxwell 16 cell DNA purification kit (Promega) on an automated DNA extraction machine (Maxwell 16 extraction system, USA). Real time PCR assay was performed on Arial Mx system (Agilent Inc, USA) using 25 μL PCR reaction mixture containing 12.5 μL Perfecta master mix low ROX kit (Quanta Bioscience Inc, USA), 1 μL of 10 μM primers, 1 μL of probes, 7.5 μL Nuclease free water (Sigma-Aldrich, USA) and 2 μL DNA template. The thermal conditions were as follows: denaturation at 95 °C for 15 min, then 30 cycles consisting of a denaturation step at 95°C for 15 s, annealing at 50°C for 15 s and extension at 70°C for 20 s.

After completion of the run, a cycle threshold (Ct) was calculated by determining the signal strength at which the fluorescence exceeded a threshold limit. This value was analyzed using the Arial Mx system software version 3.1

### Statistical analysis

Two-way ANOVA was performed using SPSS V 21 for statistical analysis of data. A p<0.05 was considered statistically significant.

### Ethical consideration

Consent was obtained from the International Foundation against Infectious Disease in Nigeria (IFAIN), Abuja to gain access to the stored *Enterobacteriaceae* obtained between 2009 and 2016. IFAIN has the written approval by the Research Ethics Committee of all the hospitals in this study.

## Results

From Sept 2008-Dec 2016, 21,000 children were screened for bacteremia with a culture positivity rate of 12.5% (2,625). Four hundred and thirteen (413) *Enterobacteriaceae* were available for analysis, comprising *Klebsiella* species 141(34.14%), *Escherichia coli* 96(23.24%), *Enterobacter* species 42 (10.16%), *Salmonella typhi* 100 (24.21%) and others 34 (8.23%) *(Serratia, Pantoea, Citrobacter, Proteus* and *Kluyvera* species).

### Prevalence of ESBLs

The overall prevalence of ESBL producers observed in this study was 160/413 (38.74%), and the prevalence of ESBL among the isolates were 65.62%, 13.75%, 13.12%, 2.5%, 4.37% and 0.62% for *Klebsiella* species, *Escherichia coli, Enterobacter* species, *Serratia* species, *pantoea* species and *Citrobacter* species respectively (Table 1).

**Table 1:** Frequency of ESBL and Non ESBL producing isolates from blood culture.

| **Isolate** | **No of isolates** | **No of ESBL producers (%)** | **No of non ESBL producers (%)** |
| --- | --- | --- | --- |
| *Escherichia coli* | 96(23.24) | 22 (13.75) | 47 (18.57) |
| *Klebsiellaspp* | 141 (34.14) | 105 (65.62) | 36 (25.53) |
| *Enterobacter spp* | 42 (10.16) | 21 (13.12) | 18 (7.11) |
| *Serratia species* | 8 (1.93) | 4 (2.5) | 4 (1.58) |
| *Pantoeaspp* | 14 (3.38) | 7 (4.37) | 7 (2.76) |
| *Citrobacters pp* | 8 (1.93) | 1 (0.62) | 7 (2.76) |
| *Proteus mirabilis* | 3(0.72) | 0(0) | 3(1.18) |
| *Kluyveraspp* | 1(0.24) | 0(0) | 1(0.39) |
| *Salmonella typhi* | 100 (24.21) | 0(0) | 100(39.52) |
| Total (%) | 413 | 160 (38.74) | 253 (61.25) |

Patient demographics associated with ESBL-producing infections are summarized in table 2.

**Table 2:** Distribution pattern of ESBL organisms by Age and Sex

|  | Escherichia coli (n=22) | Klebsiella species (n=105) | Enterobacter species (n=21) | Pantoea species (n=7) | Serratia species (n=4) | Citrobacter species (n=1) |
| --- | --- | --- | --- | --- | --- | --- |
| **Age (months)** |  |  |  |  |  |  |
| 0-1 | 6 | 53 | 9 | 4 | 2 | 0 |
| 12-Feb | 8 | 21 | 7 | 3 | 1 | 0 |
| 13-24 | 3 | 2 | 2 | 0 | 0 | 0 |
| 25-36 | 0 | 1 | 2 | 0 | 0 | 0 |
| 37-48 | 2 | 2 | 1 | 0 | 0 | 0 |
| 49-60 | 1 | 9 | 0 | 0 | 0 | 0 |
| Unknown | 1 | 17 | 0 | 0 | 1 | 1 |
| **Sex** |  |  |  |  |  |  |
| Female | 6 | 39 | 6 | 0 | 2 | 0 |
| Male | 14 | 50 | 12 | 7 | 2 | 0 |
| Unknown | 3 | 16 | 3 | 0 | 0 | 1 |

Overall, the prevalence of ESBL-producing isolates was greater among males 53.12% as compared to females 33.12% while 13.75% was unknown. ESBL-producing *Klebsiella. pneumoniae, Enterobacter* species, *Pantoea* species and *Serratia* species occurred most frequently in children ≤ 1 month of age, while *Escherichia coli* was isolated more from age range 2-12 months.

ESBL distribution in pediatric patients from Kano and FCT shows that, FCT 103/160(64.37%) had more ESBL compared to Kano 57/160 (35.62%), there was no significant difference (p=0.813; p>0.05) Fig 2.

### Antibiotic susceptibility data

Susceptibility profile of the 160 ESBL-producing isolates revealed high resistance rates for Ceftriaxone (92.3%), Aztreonam (96.8%), Cefpodoxime (96.25%), Cefotaxime (98.75%) and sulphamethoxazole-trimethoprim (90%). Over 80% of *the* isolates were resistant to Cefepime, Amoxicillin clavulanic acid and Ceftazidime (Fig 1), while 87.5 %, 90.63%, and 91.87% of the isolates were susceptible to Imipenem, Amikacin and Meropenem respectively.

**Fig 1:**
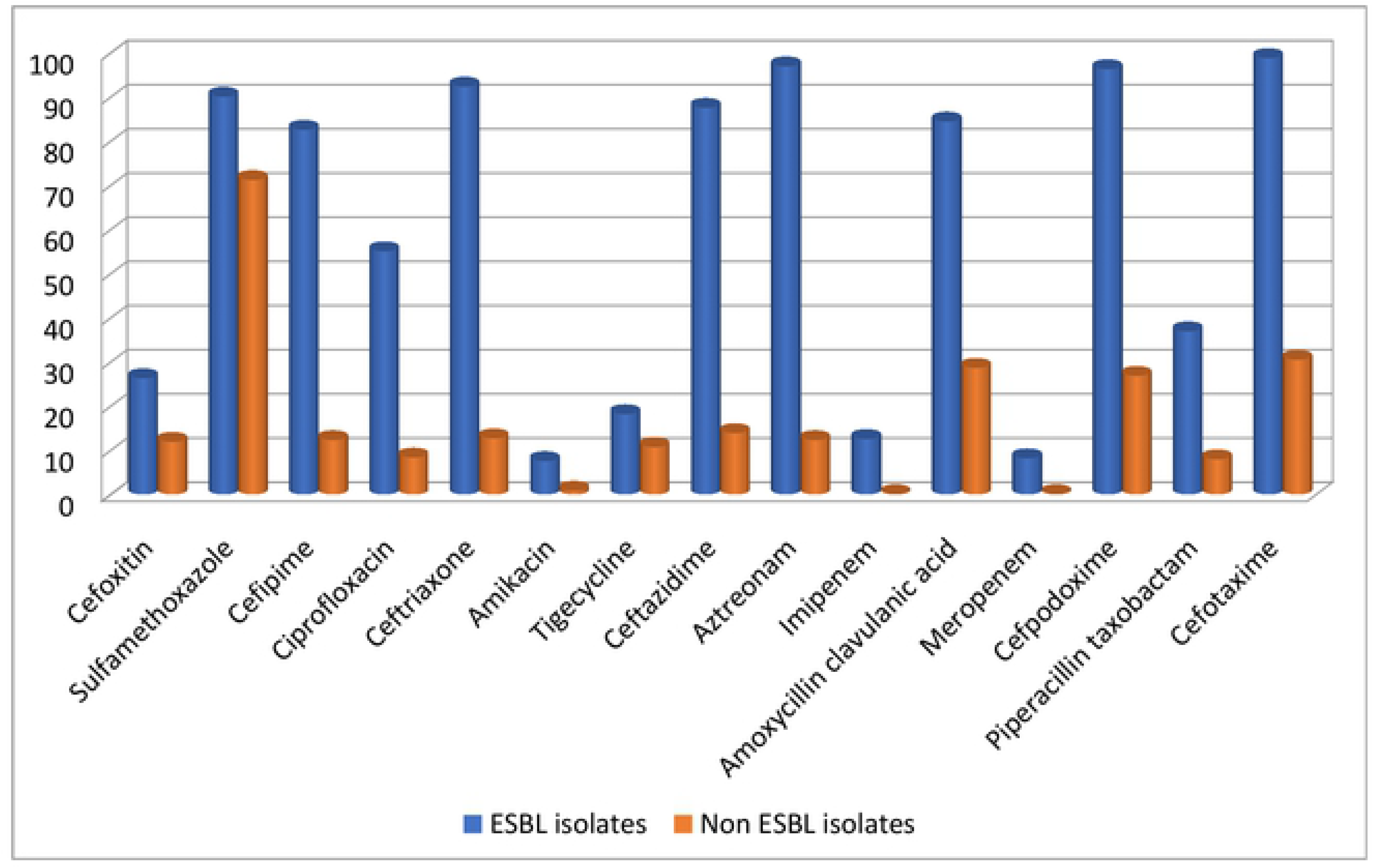
Resistance pattern of ESBL and Non ESBL producing *Enterobacteriaceae*

**Fig 2:**
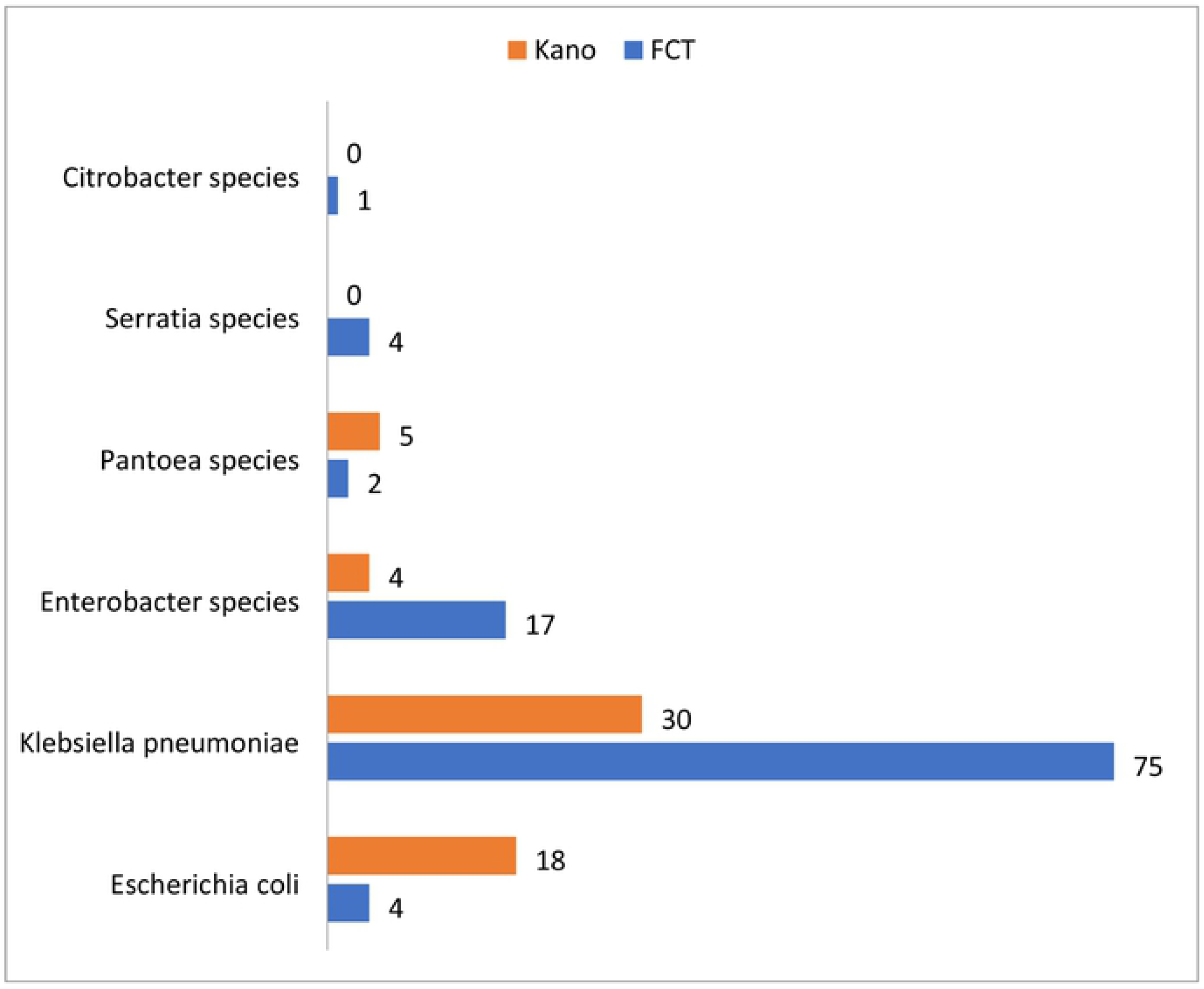
Distribution pattern of ESBL organisms from FCT and Kano.

### *bla* gene composition of ESBL-producing strains

Analysis of the phenotypically confirmed isolates revealed that 134 (62.79%) had the TEM gene, out of which 94(48.14%) were *Klebsiella pneumoniae*, 15 (25.93%) were *Escherichia coli*, 19 (22.22%) were *Enterobacter* species, 2 *Serratia* species, 3 *Pantoea* species and 1 strain of *Citrobacter* species. Also, 133 (34.88%) of the total isolates had the CTX-M gene with 98 (53.33%) being *Klebsiella pneumoniae*, 13 (6.67%) *Escherichia coli*, 5 (33.33%), 16 *Enterobacter* species, 3 were *Serratia* species, 2 *Pantoea* species and one isolate of *Citrobacter* species and 66.25% (106/160) of the total isolates had the SHV gene out of which 90/106 (66.67%) were *Klebsiella pneumoniae*, 3 were *Escherichia coli*, 10 *Enterobacter* species, 2 *Pantoea* species, and1 *Serratia* species (Fig 3). Some of the isolates expressed multiple occurrences of genes, the coexistence of blaCTX-M, blaTEM and blaSHV was seen in 94 of the isolates, while blaCTX-M and blaTEM co-existed in 118 of the isolates, blaTEM and blaSHV in 97 of the isolates while blaCTX-M and blaSHV in 100 of the isolates. Two *Escherichia coli* isolates expressed ESBL phenotype but no blaTEM, blaSHV or blaCTX-M was detected by PCR.

**Fig 3:**
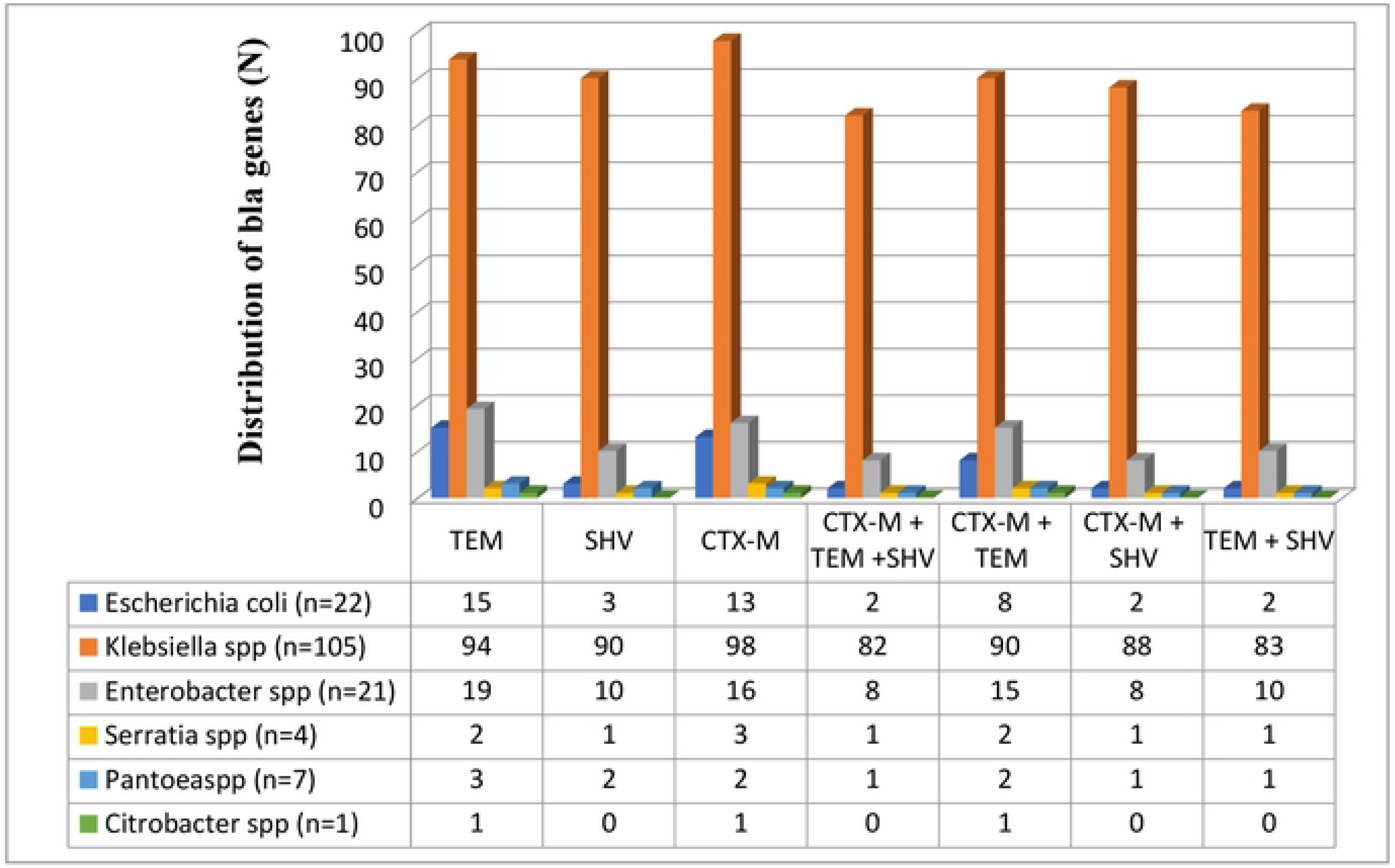
Prevalence of ESBL genes among *Enterobacteriaceae* isolates

## Discussion

ESBLs have become a widespread serious problem and these enzymes are becoming increasingly expressed by many strains of pathogenic bacteria [23]. The rate of ESBL-producing *Enterobacteriacea* in our study was 38.74% and the isolates were multi drug resistant. ESBL producing isolates showed a higher degree of antimicrobial resistance as compared to non-ESBL producers. Carbapenem and aminoglycosides were shown to be the most effective antimicrobials for ESBL and Non ESBL isolates. In comparison with non ESBL, there was a significant difference in the antibiotic resistance pattern (P= 0.0004; p>0.05).

From our study, it was observed that there were more ESBL isolates from FCT compared to Kano; this probably might be due to the affordability and access to third generation cephalosporins in FCT. Few studies have investigated the prevalence and genetic characteristics of ESBL producing *Enterobacteriaceae* from blood stream infections of pediatric patients in Nigeria. Kasap et al. 2010 from Southwestern Nigeria reported the isolation of SHV 12 from the blood of a 2-year-old girl [24], and Aibinu et al., conducted a study in 2003 in Lagos and found that eight out of 40 *Enterobacter* isolates (20%) investigated were ESBL producers [25]. The limitations in comparison with their studies were the sample size and study period. Their study covered between a period of one month and nine months, while ours had a bigger sample size and a study period of 8 years (September 2008 to December 2016)

In 2012, Alo et al., detected 80% of ESBL production among *Klebsiella pneumoniae* and *Escherichia coli* strains isolated from blood samples of hospitalized patients in Ebonyi State University Teaching Hospital [26]. Also, a study by Adeyankinnu et al., 2014 reported an ESBL prevalence of 26.4% for all isolates tested, with *E. coli* having a greater proportion [27]. Their study was restricted to detect the presence of ESBL in *Escherichia coli* and *Klebsiella pneumoniae* isolates by phenotypic means only.

The data from our study have demonstrated that there is a high prevalence of blaTEM, blaCTX-M, and blaSHV ESBL genes in *Enterobacteriaceae* isolates. A study from South Eastern Nigeria reported blaCTX-M-15 genes from urine, vaginal and wound swabs of out-patients younger than 30 years [28]. Two studies conducted in Western Nigeria by Olowe *et al*., [29] from blood, wound, HVS, and sputum samples and Raji *et al*., [30] from blood, Urine and wound samples revealed the isolates harbored blaCTX-M-1 and blaCTX-M-15 genes respectively.

According to our study, blaTEM and blaCTX-M type were the most prevalent ESBL encoding genes, detected in 83% of the ESBL-producing *Enterobacteriaceae* and the majority were found in *Klebsiella* isolates, in contrast to our study, Mohammed *et al*., from North Eastern Nigeria reported blaSHV (36.4%) and blaTEM (31.4%) to be the most prevalent [31], although there is a variation in the study design in that our study source of sample was only blood while Mohammed *et al*., used various specimen [31]. The detection of CTX-M, TEM and SHV genes by molecular techniques in ESBL producing bacteria can supply useful data about their epidemiology, association with epidemic clones and risk factors associated with these infections.

The results from our study however, reflect the global trend toward a pandemic spread of CTX-M-type ESBLs in various *Enterobacteriaceae*. These findings agree with other contemporary studies from around the world that show that ESBL genes of the CTX-M are dominant in Tanzania, Burkina Faso, Texas, Spain, Brazil, Latin America, [32–37]. In similarity to our findings, Ahmed et al. [38] reported that blaTEM was the most frequent β-lactamase-encoding gene in Egypt. In the present study, it was observed that there were multiple occurrences of genes in some of the isolates, this finding is similar to a study in Peru by Garcia *et al*., where majority (57.3%) of the ESBL strains harbored 2 or more ESBL genes [39] while in Turkey, Bali *et al*., observed that about 19.2% ESBL isolates carried more than one type of beta lactamases genes [40]. There were 2 isolates that had none of the genes tested for in them from this study, this may be due to the presence of ESBL genes other than TEM, SHV and CTX-M.

In conclusion, our findings suggest a high prevalence of ESBL resistance to commonly-used antibiotics in *Enterobacteriaceae* bacteremia in children in this study. Further studies on the transmission dynamics of resistance genes could help in the control of ESBL resistance in these settings.

## References

1. Obaro, S., Lawson, L., Essen, U., Ibrahim, K., Brooks, K., Otuneye, A. Adegbola, R. Community acquired bacteremia in young children from central Nigeria--a pilot study. BMC Infectious Diseases, 2011; 11, 137.

2. Livermore DM, et al. CTX-M: changing the face of ESBLs in Europe. J. Antimicrob. Chemother. 2007; 59:165–174.

3. J. D. Pitout and K. B. Laupland, “Extended-spectrum β-lactamase-producing *Enterobacteriaceae*: an emerging public-health concern,” The Lancet Infectious Diseases, vol. 8, no. 3, 2008, 159–166, 2008.

4. Wilke MS, Lovering AL, Strynadka NCJ. Beta-lactam antibiotic resistance: a current structural perspective. Curr Opin Microbiol. 2005; 8: 525–33.

5. Livermore DM, Woodford N. The beta-lactamase threat in *Enterobacteriaceae*, Pseudomonas and Acinetobacter. Trends Microbiol. 2006; 14: 413–20.

6. World Health Organization (WHO): Antimicrobial Resistance Global Report on Surveillance. 2014

7. Jacoby, G.A. and Munoz-Price, L.S. The new beta-lactamases. N Engl J Med. 2005; 352: 380–391

8. Demir S, Soysal A, Bakir M, Kaufmann ME, Yagci A. Extendedspectrum beta-lactamase-producing *Klebsiella pneumoniae* in paediatric wards: a nested case-control study. J. Paediatr. Child Health 2008; 44:548–553.

9. Dhanji H, et al. Real-time PCR for detection of the O25b-ST131 clone of *Escherichia coli* and its CTX-M-15-like extended-spectrum betalactamases. Int. J. Antimicrob. Agents 2010; 36:355–358.

10. Hawkey PM. Prevalence and clonality of extended-spectrum betalactamases in Asia. Clin. Microbiol. Infect. 2008;159–165.

11. Peirano G, et al. High prevalence of ST131 isolates producing CTX-M-15 and CTX-M-14 among extended-spectrum-beta-lactamase producing *Escherichia coli* isolates from Canada. Antimicrob. Agents Chemother. 2010; 54:1327–1330.

12. Tian GB, et al. Detection of clinically important beta-lactamases in commensal *Escherichia coli* of human and swine origin in western China. J. Med. Microbiol. 2012; 61:233–238.

13. Meeta S, Sati P, Preeti S. Prevalence and antibiogram of extended spectrum β-lactamase (ESBL) producing Gram negative bacilli and further molecular characterization of ESBL producing *Escherichia coli* and *KlebsiellaSpecies*. J of Clin and Diag Res. 2013;7(10):2173–77.

14. Egbebi AO, Famurewa O. Prevalence of extended spectrum beta lactamases production among Klebsiellaisolates in some parts of South West Nigeria. J Microbiol Biotech Res. 2011;1(2):64–68.

15. Aibinu IE, Ohaegbulam VC, Adenipekun EA, Ogunsola FT, Odugbemi TO, Mee BJ. Extended spectrum beta lactamase enzymes in clinical isolates of *Enterobacter* Species from Lagos, Nigeria. J ClinMicrobiol. 2003;41:2197–200.

16. Aboderin OA, Adefehinti O, Odetoyin BW, Olotu AA, Okeke IN, Adeodu OO. Prolonged febrile illness due to CTX-M-15 extended spectrum beta lactamase producing *Klebsiella pneumoniae* infection in Nigeria. Afr J Lab Med. 2012;1(1):1–4.

17. Yusha’u MM, Aliyu HM, Kumurya AS, Suleiman L. Prevalence of extended spectrum beta lactamases among *Enterobacteriaceae* in Murtala Muhammad Specialist Hospital, Kano, Nigeria. Bajopas. 2010;3(1):169–77.

18. Olowe OA, Aboderin BW. Detection of extended spectrum beta lactamase producing strains of *Escherichia coli* and *Klebsiella* Species in a tertiary health centre in Ogun state. Int J of Trop Med. 2010;5(3):62–4.

19. Olusegun O. Soge, Anne Marie Queenan, Kayode K. Ojo, Bolanle A. Adeniyi, Marilyn C. Roberts; CTX-M-15 extended-spectrum β-lactamase from Nigerian *Klebsiella pneumoniae*, Journal of Antimicrobial Chemotherapy, Volume 57, Issue 1, 1 January 2006, Pages 24–30,

20. Obaro, S. K., Hassan-Hanga, Fey, P. D et al. Salmonella Bacteremia Among Children in Central and Northwest Nigeria, 2008-2015. Clinical infectious diseases: an official publication of the Infectious Diseases Society of America, 2015;61 Suppl 4(Suppl 4), S325–31.

21. CLSI. Performance standards for antimicrobial susceptibility testing, 27^th^ ed. CLSI supplement M100. C Wayne, PA; Clinical and Laboratory Standards Institute; 2017.

22. Roschanski N, Fischer J, Guerra B, Roesler U. Development of a Multiplex Real-Time PCR for the Rapid Detection of the Predominant Beta-Lactamase Genes CTX-M, SHV, TEM and CIT-Type AmpCs in *Enterobacteriaceae*. PLoS ONE 2014;9(7): e100956.

23. Sibhghatulla Shaikh, Jamale Fatima, Shazi Shakil, Syed Mohd. Danish Rizvi, Mohammad Amjad Kamal. Antibiotic resistance and extended spectrum beta-lactamases: Types, epidemiology and treatment Saudi J Biol Sci. 2015 Jan; 22(1): 90–101

24. Kasap, M., Fashae, K., Torol, S., Kolayli, F., Budak, F., & Vahaboglu, H. Characterization of ESBL (SHV-12) producing clinical isolate of Enterobacter aerogenes from a tertiary care hospital in Nigeria. Annals of clinical microbiology and antimicrobials, 2010;9, 1.

25. Aibinu, I. E., Ohaegbulam, V. C., Adenipekun, E. A., Ogunsola, F. T., Odugbemi, T. O., & Mee, B. J.. Extended-spectrum beta-lactamase enzymes in clinical isolates of Enterobacter species from Lagos, Nigeria. Journal of clinical microbiology, 2003;41(5), 2197–200.

26. Alo, M. N, Anyim C., J. C. Igwe and Elom, M. Presence of extended spectrum β-lactamase (ESBL) E. coli and K. pneumonia isolated from blood cultures of hospitalized patients. Advances in Applied Science Research 2012; 3 (2):821–825.

27. Adeyankinnu, F. A., Motayo, B. O., Akinduti, A., Akinbo, J., Ogiogwa, J. I., Aboderin, B. W., and Agunlejika, R. A. A Multicenter Study of Beta-Lactamase Resistant Escherichia coli and Klebsiella pneumoniae Reveals High Level Chromosome Mediated Extended Spectrum β Lactamase Resistance in Ogun State, Nigeria. Interdisciplinary perspectives on infectious diseases, 2014;896.

28. Iroha I R, Esimone C O, Neumann S et al. “First description of *Escherichia coli* producing CTX-M-15-extended spectrum betalactamse (ESBL) in out-patients from South Eastern Nigeria. Annals of Clinical Microbiology and Antimicrobials 2012:11–19.

29. Olowe O A, Oladipo G O, Makanjuola O A, and Olaitan J O. “Prevalence of Extended Spectrum Betalactamases (ESBLs) Carrying Genes in Klebsiellaspeciesfrom Clinical Samples at Ile-Ife, South Western Nigeria,” International Journal of Pharma Medicine and Biological Sciences, Vol. 1, No. 2, pp. 129–138, October 2012.

30. Raji, M A, Jamal W, Ojime O and Rotimi V O. “Sequence Analysis of genes Mediating Extended Spectrum Beta Lactamase (ESBL) production in isolates of *Enterobacteriaceaein* a Lagos Teaching Hospital, Nigeria. BMC Infectious Diseases 2015; 15:259.

31. Mohammed Y, Gadzama GB, Zailani SB, Aboderin AO. Characterization of Extended-Spectrum Beta-lactamase from *Escherichia coli* and *KlebsiellaSpecies* from North Eastern Nigeria. Journal of Clinical and Diagnostic Research: JCDR. 2016;10(2): DC07–DC10.

32. Moremi N, Claus H, Vogel U, Mshana SE. Faecal carriage of CTX-M extended-spectrum beta-lactamase-producing *Enterobacteriaceae* among street children dwelling in Mwanza city, Tanzania. PLoS ONE (2017) 12(9): e0184592

33. Ouedraogo, Abdoul-Salam et al. “High Prevalence of Extended-Spectrum SS-Lactamase Producing *Enterobacteriaceae* among Clinical Isolates in Burkina Fas.” BMC Infectious Diseases 16 (2016): 326.

34. Chandramohan, Lakshmi, and Paula A. Revell. “Prevalence and Molecular Characterization of Extended-Spectrum-B-Lactamase-Producing *Enterobacteriaceae* in a Pediatric Patient Population.” Antimicrobial Agents and Chemotherapy 56.9 (2012): 4765–4770.

35. Díaz, Miguel A. et al. “Diversity of *Escherichia Coli* Strains Producing Extended-Spectrum B-Lactamases in Spain: Second Nationwide Study.” Journal of Clinical Microbiology 48.8 (2010): 2840–2845.

36. NogueiraKeite da Silva, Conte Danieli, Maia Fernanda Valverde, Dalla-Costa Libera Maria. Distribution of extended-spectrum β-lactamase types in a Brazilian tertiary hospital. Rev. Soc. Bras. Med. Trop. 2015 Apr 48(2): 162–169

37. Pallecchi L, Bartoloni A, Fiorelli C, et al. Rapid Dissemination and Diversity of CTX-M Extended-Spectrum β-Lactamase Genes in Commensal *Escherichia coli* Isolates from Healthy Children from Low-Resource Settings in Latin America. Antimicrobial Agents and Chemotherapy. 2007;51(8):2720–272

38. Ahmed SH, Daef EA, Badary MS, Mahmoud MA, Abd-Elsayed AA. Nosocomial blood stream infectionin intensive care units at Assiut University Hospitals (Upper Egypt) with special reference to extendedspectrum beta-lactamase producing organisms. BMC Res Notes. 2009; 2: 76.

39. García C, Astocondor L, Rojo-Bezares B, Jacobs J, Sáenz Y. Molecular Characterization of Extended-Spectrum β-Lactamase-Producer *Klebsiella pneumoniae* Isolates Causing Neonatal Sepsis in Peru. The American Journal of Tropical Medicine and Hygiene. 2016;94(2):285–288.

40. Bali BE, Acik L, Sultan N. Phenotypic and molecular characterization of SHV, TEM, CTX-M and extended spectrum beta lactamases produced by *Escherichia coli, Acinobacter baumannii* and *Klebsiella* isolates in a Turkish Hospital. Afr J of Res. 2010;4(8):650–54.

